# Spectroscopic and morphological study of sinusoidal mechanical vibration exposed human RBC in vitro

**DOI:** 10.1101/2025.06.08.658542

**Authors:** Vijay Ghodake, Sarika Hinge, Arun Banpurkar, Gauri Kulkarni

## Abstract

Humans undergoing mechanical vibration of frequency ranging from mHz to a few kHz in daily life. Human body contains about 60% of water. Blood is one of the most important body fluids and RBC are functional constituents of blood. RBC, due to their deformable nature, can undergo physical deformation or membrane damage under the influence of external mechanical vibrations over extended periods. In the present work, we studied in vitro the effect of sinusoidal mechanical vibration of 2 mm amplitude and frequency ranging from 5 to 100 Hz for 10 min duration. Physicochemical properties of controlled and mechanically vibrated RBC were examined using UV-visible, Fourier Transform Infrared (FTIR), Raman spectroscopy, and morphology was examined by Scanning Electron Microscopy (SEM). Results showed that mechanical vibrations affect oxygenation level at the Q band (543 nm and 578 nm), and variation in transmission intensity is observed in the Amide region. RBC showed a change in morphology, where the elongation index increases along with vibration frequency. Current findings are useful to various stages of blood management and supply chain like collection, storage, transportation, handling, and donation.

**Key points:**

- Effect of mechanical vibration on RBC highlights the importance of blood transportation and handling related issues.
- Spectroscopic and Morphological analysis revealed the modification in RBC due to sinusoidal mechanical vibration.

## Introduction

Daily physical activities like running, walking, playing, driving, travelling cause excessive mechanical stress viz. compression, tension, and shear stress on human body.^**1-3**^ During road transportion, human body experience mechanical vibrations in the range of 1 to 500 Hz ^**4**^ which indirectly affects the organs, tissues and soft matter in human body. As a result, damage to the cytoskeleton, proteins, cells, and tissues occur which alters cell cycle, its metabolism and blood supply to various vital organs. ^5,6^ Mechanical vibrations can be destructive or constructive in nature and generate abiotic stresses on the body cells^**7-8**^ which depends on its waveform, frequency, amplitude, and exposure time. In some cases, vibrational frequencies affect the bone density, nerve, muscle, vision and skin microcirculation. ^**9-13**^ High frequency vibrations can also create multifracture damage to the bone. ^**14**^ However, whole human body treated with external vibrations during physical workout showed better performance in bone strength and bone mass in adult persons, astronauts, etc.^15^ Vibrational frequency is used in physiotherapy for treatments of joints and tissues, hemodialysis, wound healing in diabetic foot ulcers, ^**16**^ morphogenesis of neuronal cells and fibroblast. ^**17**,**18**^ In recent years, it is shown that the mechanical vibration affects the blood circulation, blood constituents and it activates. ^**19**^ Changes in the physicochemical characteristics of RBC and its biological pathways might result in circulation disorders, and cardiovascular problem due to mechanical vibrations. ^**20**,**21**^ About 45% blood volume is occupied by RBC and it is a major constituent of whole blood mainly used as a gas transporter and drug carrier. Blood components like RBC, lymphocytes, platelets and blood plasma get affected due to mechanical vibration ^**22**,**23**^ which influences the viscosity of blood to avoid aggregation ^**24**^ and increases the erythrocyte volume fraction (EVF) of arm hand and Hb in index fingers.^**25**^ Other studies have also shown that vibrations and mechanical shock can increase the hemolysis of RBC. External local vibration therapy delivered to impaired part of the human body increases blood flow without any RBC aggregation. However, most of the vibrations show adverse effect on peroxidase activity of hemoglobin at low frequency vibrations within the range of 8 to 32 Hz. ^**26**^ Earlier, it was reported that road transportation stress on RBC causes osmotic fragility of RBC, hemoglobin level and blood rheology. ^**27**^

Although the effects of mechanical vibrations on biological systems have been documented, it is not known how mechanical vibrations affect morphological and biophysical properties of RBC. In the present study, RBC were exposed to 5 Hz,10 Hz, 20 Hz, 30 Hz, 40 Hz, 50 Hz, 60 Hz, 70 Hz, and 100 Hz sinusoidal vibrational frequency. Comparison between vibrated and non-vibrated RBC was done on the basis of UV-visible spectroscopy, FTIR spectroscopy, Raman spectroscopy and Scanning Electron Microscopy.

## Materials and Methods

### Sample preparation

Blood sample (10 ml) was collected from venous of healthy human by in ethylenediamine tetraacetic acid ∼EDTA, (3g/l) and all methods were performed with the approval and following the relevant guidelines and regulations set by the Institute Ethical Committee (Ref. No. SPPU/IEC/2021/129). Collected blood was stored at 4 °C and used for the experiment within 48 hr. RBC were extracted from whole blood at 3000 revolutions per minute (RPM) for 10 minutes at 25 °C, using centrifugation process. Blood components like RBC, WBC, plasma were separated relative to density gradient. Isolated RBC were washed thrice by using 0.9% saline and used for experiments. All experiments were conducted three times using pure RBC samples obtained from adult, healthy volunteers (N = 5).

### Experimental setup for mechanical vibration

A vibration test system (Sine force – peak 100 kgf, frequency range 5 Hz to 4500 Hz, maximum acceleration, velocity, and displacement are 65 g, 1.6 m/s, and pk-pk 25 mm, respectively) using an electrodynamic shaker from Spectral Dynamics, USA was used to induce mechanical vibrations of various frequencies to the sample holder the sample. The schematic of the mechanical vibration system is shown in Figure 1.

**Figure 1.**
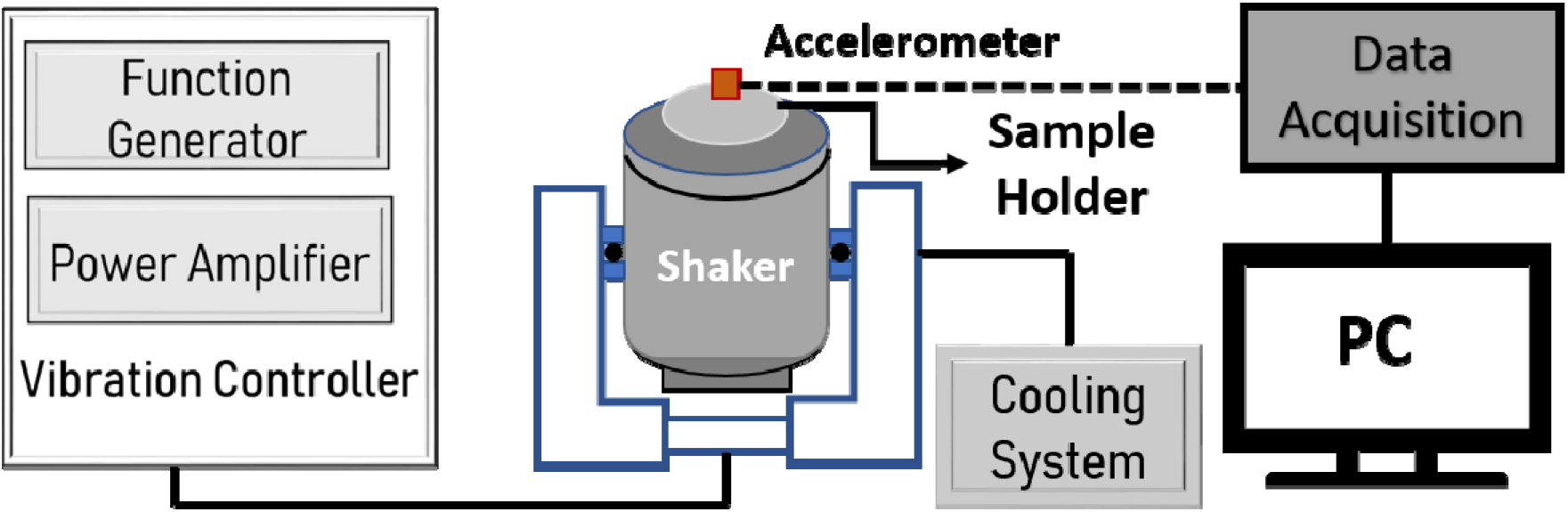
Schematic of the experimental setup for inducing mechanical vibration system.

The driver signal from the function generator is input to the amplifier. The Amplifier converts low voltage and mA currents from the controller to higher voltages and currents require for the driving shaker unit. The moving element (armature) of the shaker is suspended in a strong magnetic field. The current output from the amplifier flows through the wire in a magnetic field. The suspended moving element is restrained and allowed to move up and down. Movement of sample table is sensed by the piezoelectric accelerometer. The conditioned signal is fed into the closed control loop from the charge amplifier to a pre-defined reference level. The control loop adjusts the drive signal to the amplifier in order to maintain the reference level. The motion of the table follows the characteristics of the current flowing through a coil produces sinusoidal vibration motion of the sample holder.

To examine the vibration effect on blood samples (0.05 ml) in vial was fixed vertically on vibration platform of electrodynamic shaker. RBC samples were vibrated at different frequencies viz. 5, 10, 20, 30, 40, 50, 60, 70, and 100 Hz with constant amplitude of 2 mm and 10 minute exposure.

### Experimental procedure for UV-visible, FTIR and Raman spectroscopy

Effect of mechanical vibrations on RBC was characterized by using UV-visible (JASCO-670, Japan), FTIR (JASCO-6100, Japan) and Raman spectroscopy (inVia™ Renishaw). UV-visible spectroscopy of RBC was studied by diluting the fluids in the ratio of 1:100 within the spectral range 200 - 800 nm. For FTIR spectra, 15 µl of RBC added into KBr powder and dried at room temperature and measurement were recorded in 400 – 4000 cm^-1^. To obtained the Raman spectra, 10 µl of RBC solution was dried on glass slide and spectra were monitored in the spectral range of 400 -1800 cm^-1^.

### Experimental procedure for Scanning Electron Microscopy of RBC

Isolated RBC (2 ml) were added in a vial containing buffer solution (0.4 ml of 2% glutaraldehyde and 19.6 ml saline). These samples were slowly shaken for 2 to 3 minutes and kept in refridgerator 10-12 hr at temperature 4 °C. Again 0.5 ml saline was added to each vial, then these vials were centrifuged and supernatant plasma was removed with micropipette. The ethanol treatment was given to the leftover red blood cells. For ethanol treatment, the solution was prepared by mixture of ethanol and saline with concentration 10, 20, 30, 40, 50, 60, 70, 80, 90, and 100 % by using the standard protocol (Add 0.2 ml ethanol to 1.8 ml saline, the total solution is 10%. Add 0.4 ml ethanol to 1.6 ml saline the total solution is 20%, similar way 30%, 40%, 50%, 60%, 70%, 80%,90% and 100% solution were prepared). Three glass slides for each sample were prepared and RBC smeared uniformly on the glass substrate. Total thirty slides were prepared and kept in vacuum for whole night. SEM for control and vibrated samples were performed for x3000 magnification.

## Results

UV-visible spectra, FTIR spectra, Raman spectra and Scanning Electron Microscopy measured for control RBC and mechanically vibrated RBC.

### UV-visible spectroscopy

The absorption spectra measured from the vibrated RBC at different frequencies and non-vibrated (control) isolated RBC from whole blood is shown in Figure 2. Absorption from vibrated and non-vibrated RBC were seen at 281 nm, 346 nm, and 418 nm, 543 nm, and 578 nm.

**Figure 2.**
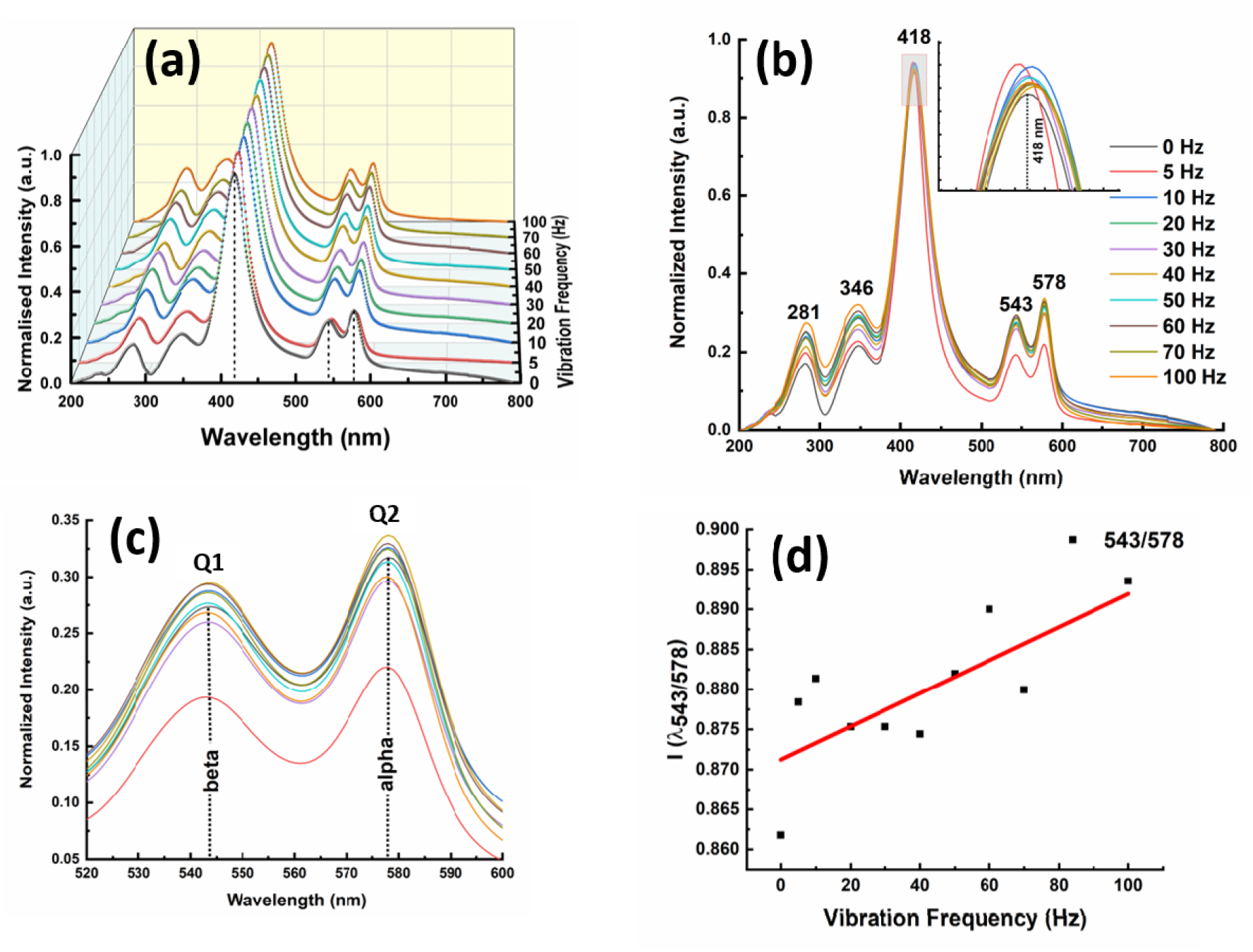
Changes in the absorption spectra of **(a)** RBC vibrated at 5, 10, 20, 30, 40, 50, 60, 70 and 100 Hz with amplitude of 2 mm for 10 minute (b) cumulative absorption spectra of RBC, inset shows intensity of 418 nm sorat band. (c) magnified view of the Q-band (d) plot shows ratio of Q1/Q2 versus vibrated frequency.

The absorbance at 281 nm and 346 nm is a determinant indicator of protein content in RBC. The peak at obtained at 281 nm is associated with π-π^*^ transition and peak at 346 nm showed signature of aromatic amino residues (Tryptophan, Tyrosine, and Phenelanine). Absorption at haem group of vibartaed and non-vibrated RBC is seen at 418 nm. The alteration in intensity and wavelength is related to haem group, and its interaction with polypeptide bonds surrounded by Fe group in porphyrin ring. However, Q band (alpha and beta) showed changes in ligand-porphyrin which is associated with oxygen binding and transport. Absorption peak related to oxygenation Q1(alpha) ∼543 nm and Q2 (beta) ∼578 nm showed increasing trend with increase in the vibrational frequencies from 5, 10, 20, 30, 40, 50, 60, 70, and 100 Hz as compared to control sample. Variation in intensity at Q1 and Q2 bands was observed for vibrated RBC which led to change in the oxygenation state.

### FTIR Spectroscopy

The FTIR spectra of control and vibrated RBC were measured at 10, 30, 50, 70 and 100 Hz with amplitude of 2 mm at 10-minute duration.

FTIR spectra of control and vibrated RBC show peaks at 1551-1558 cm^-1^ which corresponds to Amide II (C=O stretching of proteins), 1632-1659 cm^-1^ corresponds to Amide I (mainly C=O stretching of proteins) and 3357 - 3443 cm^-1^ corresponds to Amide A (N-H stretching of secondary amide proteins). ^**28**^ Figure 3 shows the peaks obtained from the control and vibrated RBC. FTIR spectra of vibrated RBC shows variation in the transmission intensity and changes in the functional group particularly amide band. Alterations in Amide groups of RBC was observed due to vibrations of different frequency.

**Figure 3.**
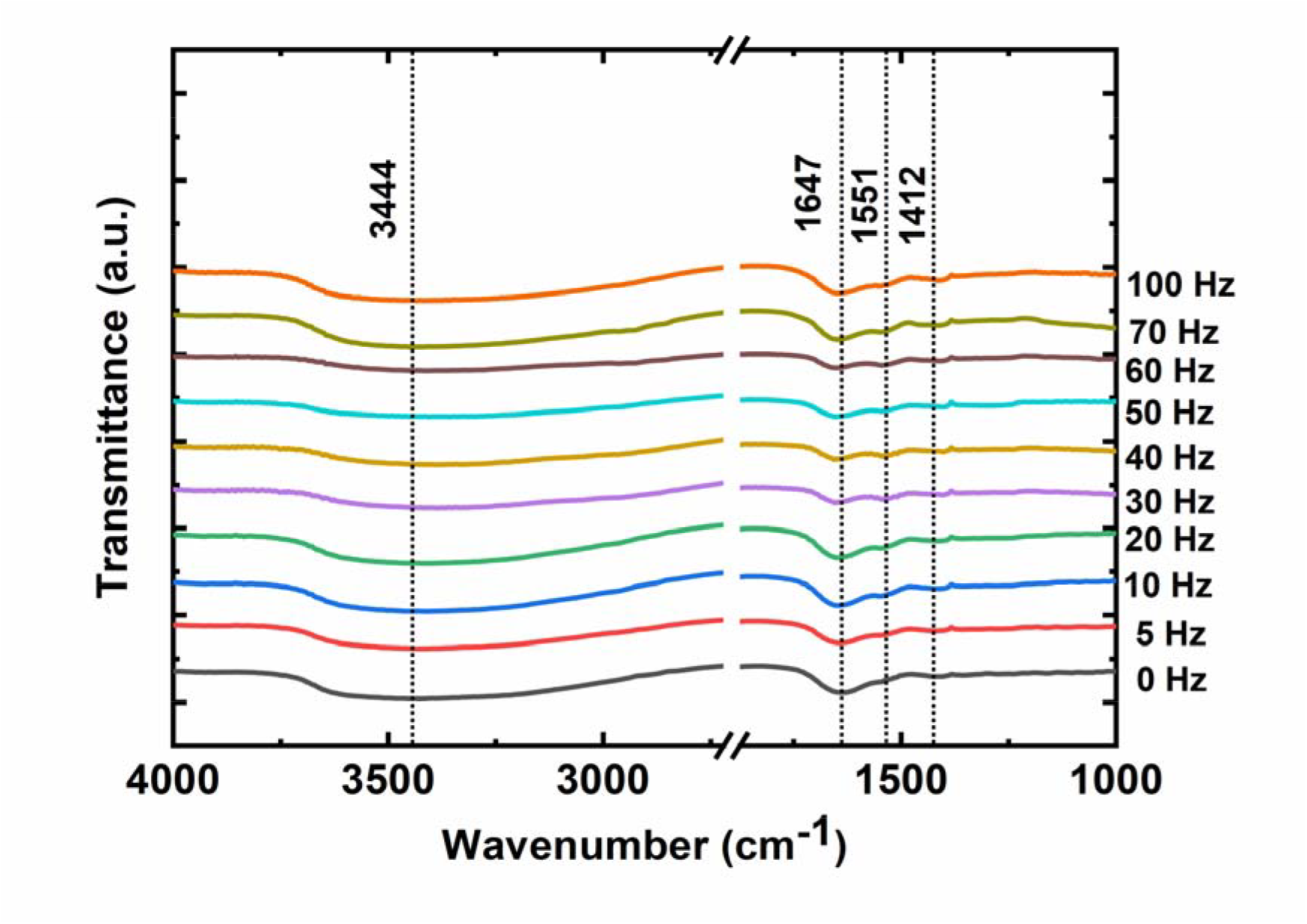
FTIR spectra of RBC vibrated at 5, 10, 20, 30, 40, 50, 60, 70, and 100 Hz with amplitude of 2 mm for 10 minute duration.

### Raman Spectroscopy

Raman spectra of RBC were recorded with laser of 785 nm wavelength (power : 3 mW) within the range of 400 cm^-1^ to 1800 cm^-1^ as shown in **Figure 4**.

**Figure 4.**
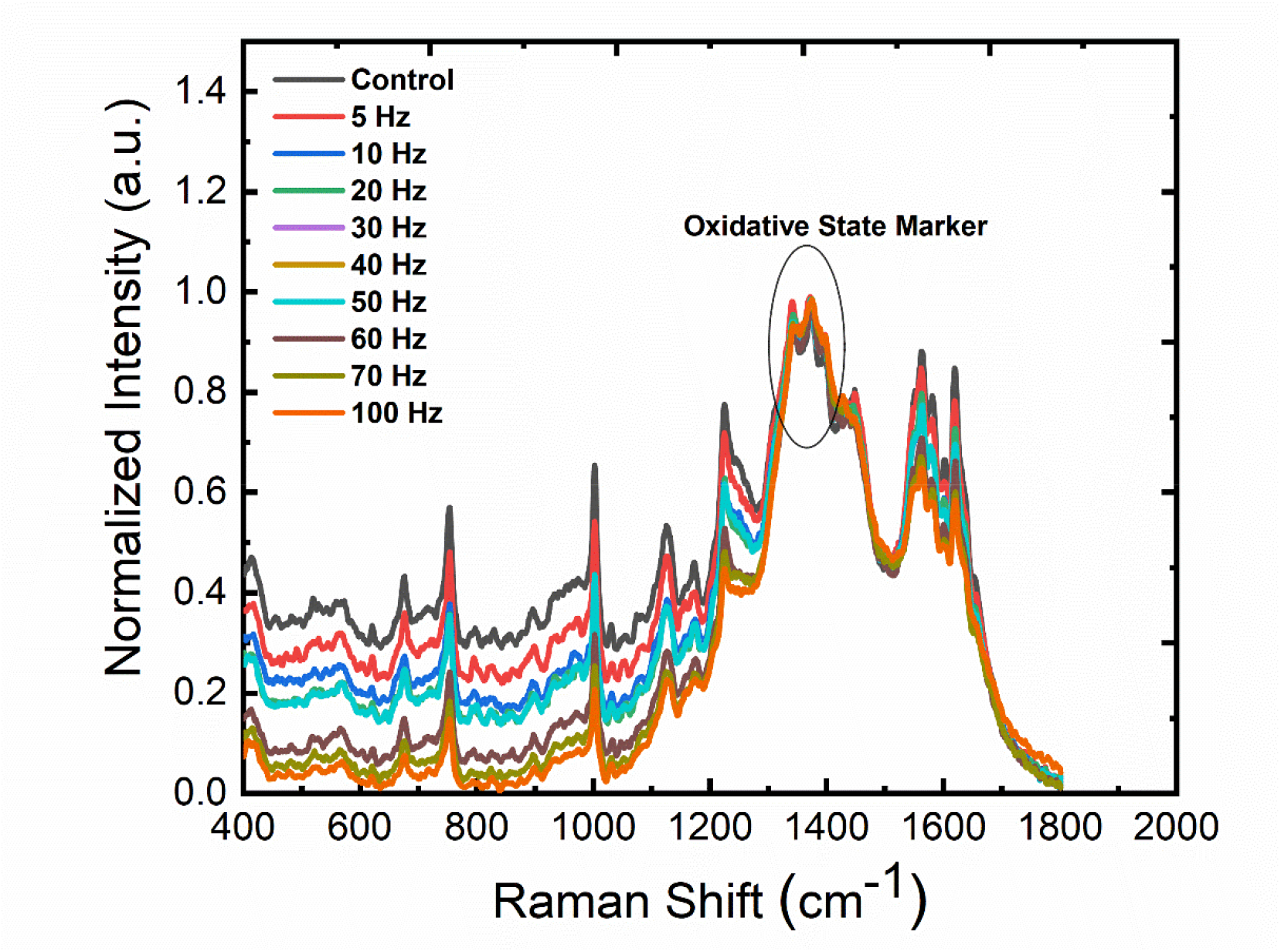
Raman spectra of RBC vibrated at 5, 10, 20, 30, 40, 50, 60, 70, and 100 Hz with amplitude of 2 mm for 10 minute duration.

Raman spectra of RBC and mechanically vibrated RBC showed the Raman peaks at 569 cm^-1^ (Fe-O_2_) stretch), 676 cm^-1^ (δ(pyrrole deformation) symmetric in plane deformation (ν_7_)), 755 cm^-1^ (ν (pyrrole breathing)stretch (ν_15_)), 793 cm^-1^ (ν(pyrrole deformation)ν_6_), 825 cm^-1^ (γ (C_m_H)out of plane deformation (γ_10_), Tyrosine), 964 cm^-1^ (C-C stretch in unordered protein),1003 cm^-1^ (ν (C_β_C_1_) asymmetric stretch (ν_45_) and Phenylalanine), 1130 cm^-1^ (C-C stretch (trans) lipids), 1175 cm^-1^ (ν(pyrrole half ring) asymmetric stretch (ν_30_)), 1224 cm^-1^ (δ(CmH) in plane deformation (ν_13_)), 1341 cm^-1^ (N (pyrrole half ring) symmetric stretch (ν_41_)), 1396 cm^-1^ (N (pyrrole quarter-ring) stretch (ν_20_)), 1450 cm^-1^ (δ(CH_2_/CH_3_)), 1564 cm^-1^ (ν(C_β_C_β_) stretch (ν_2_)),1580 cm^-1^ (ν (C_α_C_m_) asymmetric stretch (ν_37_)),1620 cm^-1^ (ν (C_a_=C_b_) of vinyl groups). ^**29**^ Normalized Raman spectra of vibrated RBC showed decrease in intensity with increase in mechanical vibration. However, oxidative state marker region (1300-1400 cm^-1^) showed slightly varition in peak position and intensity.

### Morphological study

Scanning electron microscopic images of RBC isolated from vibrated whole blood were obtained at different frequencies (Figure 5).

**Figure 5.**
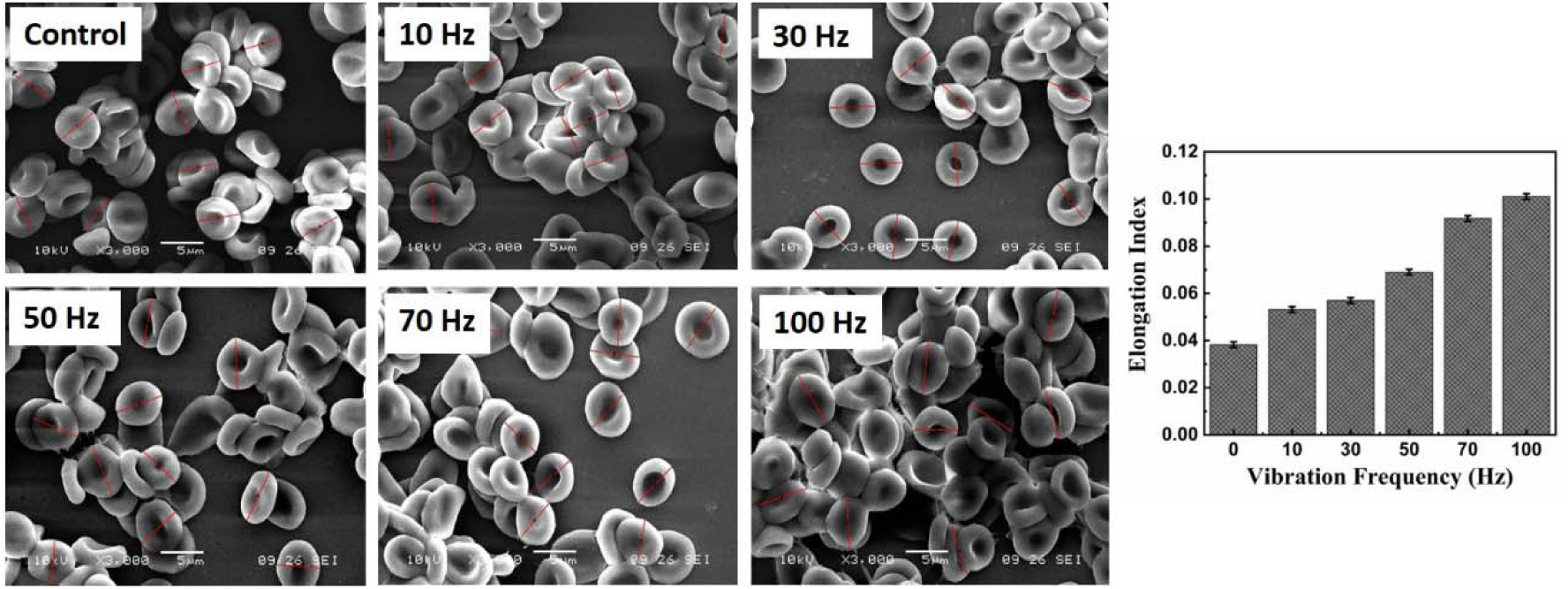
(a) Morphology of isolated RBC (Control) and vibrated RBC at different frequencies of vibrations i.e. 10, 30, 50, 70, and 100 Hz with constant amplitude (2 mm) (b) Elongation index Vs Vibrational frequency

Scanning electron microscope (JEOl JSM-6360A) was used to obtain RBC images before and after mechanical vibration. Public domain (Image J, 1.54f) software is used for data analysis. From the control, the average diameter of RBC was measured as 4.76 μm and thickness around 1.84 μm. In the case of vibrated samples, the diameter of RBC increases from 4.76 μm ± 0.08 to 5.64 ± 0.06 μm with increasing vibrating frequencies and the width of RBC decreases from 1.84 ± 0.03 μm to 1.32 ± 0.05 μm with increasing vibrating frequencies. Change in the elongation index of RBC at different vibration frequencies is seen in Figure 5(b). Scanning electron microscopic images show deformation in RBC after exposure to sinusoidal mechanical vibrations, diameter of RBC increased along with a decrease in thickness, and width. The same trend was observed in isolated RBC from vibrated whole blood at different frequencies. More details of the physicochemical properties of vibrated whole blood and its characterizations are available in the electronic supplementary note (**SP)**.

## Discussion

The advancement in the research of vibration on RBC has potential in health and risk management in blood transfusion. During blood transportation via road, many external factors affect the physiology of RBC. To maintain the efficacy and integrity of RBC, it is necessary to understand the effect of sinusoidal vibration particularly for blood transfusion.

Hemoglobin is a major part of RBC carrying and binding oxygen to an encapsulated porphyrin ring with two alpha and beta chains. Absorption at oxygen binding sites is examined at 543 nm (Q1) and 578 nm (Q2) using UV-visible spectroscopy. Mechanical vibration affects the oxygenation state of RBC which is determined by the ratio of Q1/ Q2. Change in the microenvironment of the RBC was observed with variation in vibrational frequency which led to a decrease in absorbance intensity and FWHM of the peak. For quantitative measurement and studying the effect of sinusoidal vibration on RBC, FTIR spectroscopy is used to investigate functional groups mainly associated with Amide I and Amide II.

Sinusoidal vibrations induce stress on the secondary structure present at Amide I and II region. RBC is composed of hemoglobin, spectrin network, cytoskeleton membrane proteins, and associate secondary structure viz. α-helix, β-sheet, random coil, and aggregates. The second derivative and difference in amide bands I and II provide information related to secondary structure. Molecular level investigation of RBC was performed by using Raman spectrsocsopy. Spectra shows changes in intensities in CH_3_ amino acid deformation mode, hemeagreegation, Fe – O_2_ stretch mode. Intensity of spectra increases, but no significant change in Raman shift was observed due to mechanical vibrations.

During microcirculilation, RBC deform from biconcave shape to dumbbell shape and passess through 3-5 micron size blood capillaries. Due to elastic and flexible nature, RBC regain its original shape and size. Elongation index of RBC was determined by measuring the diameter and width of RBC using public domain software (ImageJ 1.54f). Vibartion affects the morphology of RBC which causes an increase in diameter with a decrease in thickness. Overall elongation index is increased as a function of vibrational frequency. Elongation may affect the deformation property of RBC. Hence, there is more research is needed to understand the deformability of vibration exposed individual live RBC.

## Conclusions

In this work, it is seen that sinusoidal mechanical vibration affects the shape and physicochemical properties of RBC. Such change is vital in microcirculation and decides oxygenation state of RBC affecting oxygen carrying capacity. Mechanical vibrational stress beyond the natural frequency of RBC affects the morphology of RBC which led to elongation of RBC, and variation in peak intensity of Raman peak. Hence, it is advisable to avoid exposure of human body to continuos mechanical vibrations within certain range of frequency amd amplitude and type of vibrations. Vibrations may cause change in mechanical properties of RBC and affects their dynamics. This work has important significance in safety handling and risk management during blood transportation..

## Authors Information

## Authors and Affiliations

Department of Physics, Savitribai Phule Pune University

Central Water and Power Research Station, Pune

## Contributions

V.G. and S.H. contributes equally. V.G: contributes to Conceptualization, Methodology, Validation, Writing; S.H: contributes to Methodology, Investigation, Data curation, writing – draft; A.B.: Data curation, writing - review & editing, Supervision; G.K: Data curation, writing - review & editing, Supervision

## Ethics approval

The entire protocol for collection of blood sample is approved by Institute Ethical Committee (Ref. No. SPPU/IEC/2021/129).

## Consent to participate

Informed consent was obtained from all individual participants included in the study.

## Competing Interests

“The authors declare that they have no competing interests”.

## Acknowledgement

Authors are grateful to Director, Central Water and Power Research Station, Pune, India and Director of Basic Medical Science, Department of Physics, Savitribai Phule, Pune University, Pune - 411007, MH, INDIA.

## Funding

None

## Data Availability

All data generated or analyzed during this study are included in this published article.

## Notes

### Competing Interest Statement

The authors have declared no competing interest.

### Summary of Updates

There is no difference in current manuscript and previous manuscript

