## supplementary note (SP) for "Spectroscopic and morphological study of sinusoidal mechanical vibration exposed human RBC in vitro"

Physicochemical characterization of RBC and plasma isolated from vibrated whole blood blood help to understand the effect of sinusoidal mechanical vibration on blood. Previously, in main manuscript, we directly observe the vibration effect on pre - isolated red blood cell from whole blood.

**Raman Spectroscopy of vibration exposed whole blood**

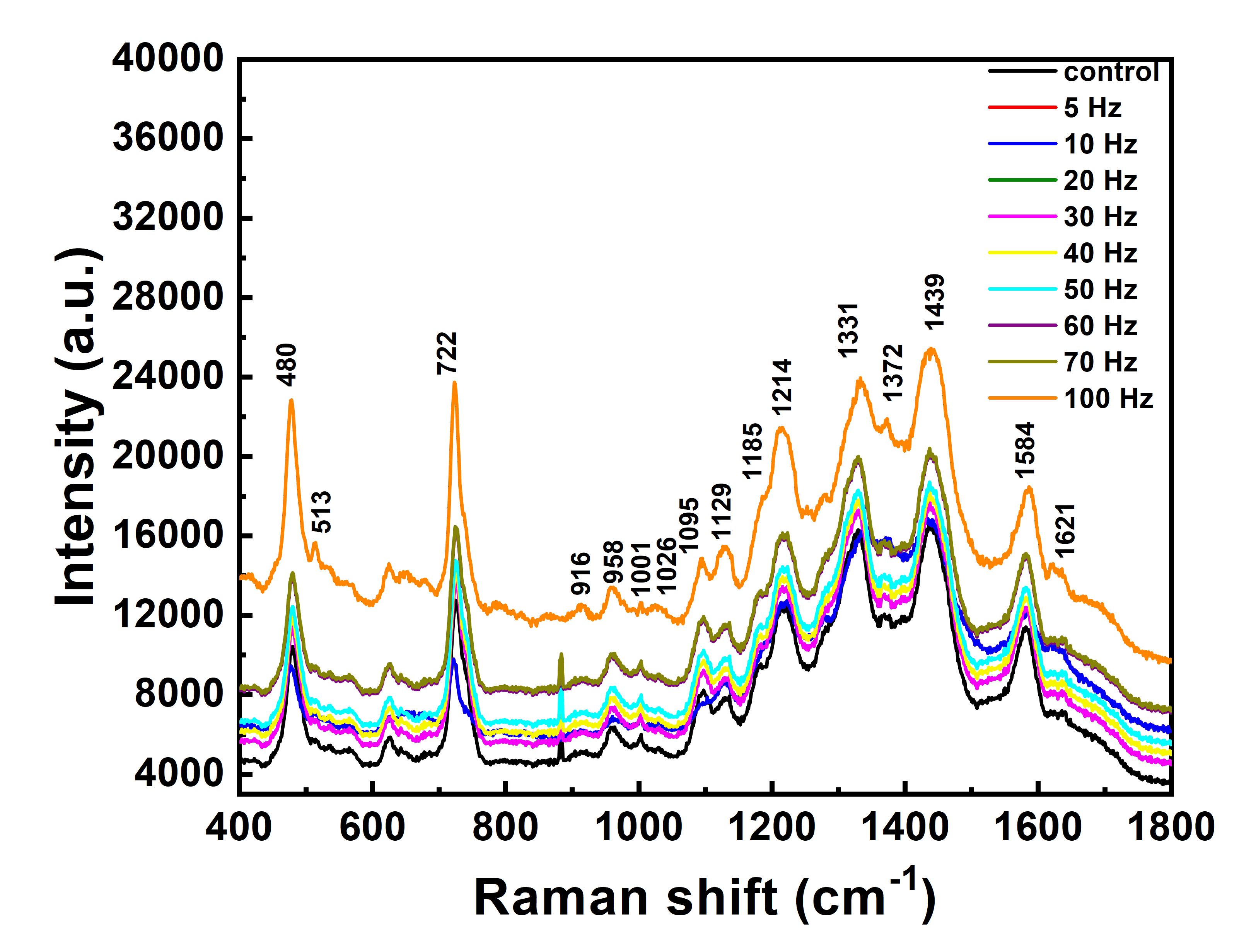

**Figure S1.** Changes in the Raman spectra of whole blood vibrated at 5, 10, 20, 30, 40, 50, 60, 70 and 100 Hz with amplitude of 2 mm at 10 minute duration.

Raman spectra of whole blood showed the Raman Peaks at 480 cm^-1^, 513 cm^-1^, 722 cm^-1^ (C-N stretch lipids), 958 cm^-1^ (C-C stretch in unordered protein), 1001 cm^-1^ (ν (C_β_C_1_) asymmetric stretch (ν_45_) and Phenelalanine), 1095 cm^-1^,( ν (C_β_C_1_)asymmetric stretch (ν_23_), C-C stretch lipids, PO_2_ - symmetric stretch, 1129 cm^-1^, (ν (pyrrole half ring) asymmetric stretch (ν_22_)), 1214 cm^-1^, (δ (CmH) in plane deformation (ν_5_)), 1331 cm^-1^,(N (pyrrole half-ring) symmetric stretch (ν_41_)), 1584 cm^-1^ (ν (C_α_C_m_) asymmetric stretch (ν_37_)) and 1621 cm^-1^(ν ( C_a_= C_b_) of vinyl groups).**^1^**

**Scanning electron microscopy of isolated RBC from vibrated whole blood**

Scanning electron microscopy of normal red blood cell showed biconcave shape.**^2^** Scanning electron microscope images of RBC in whole blood were obtained for different frequencies (**Figure S2**) . From the control the average diameter of RBC was measured as 4.76 μm and width around 1.84. In case of vibrated samples, the diameter of RBC increases from from 4.76 μm ± 0.08 to 5.64 ± 0.06 μm with increasing vibrating frequencies and width of RBC decreases from 1.84 ± 0.03 μm to 1.32 ± 0.05 μm with increasing vibrating frequencies.

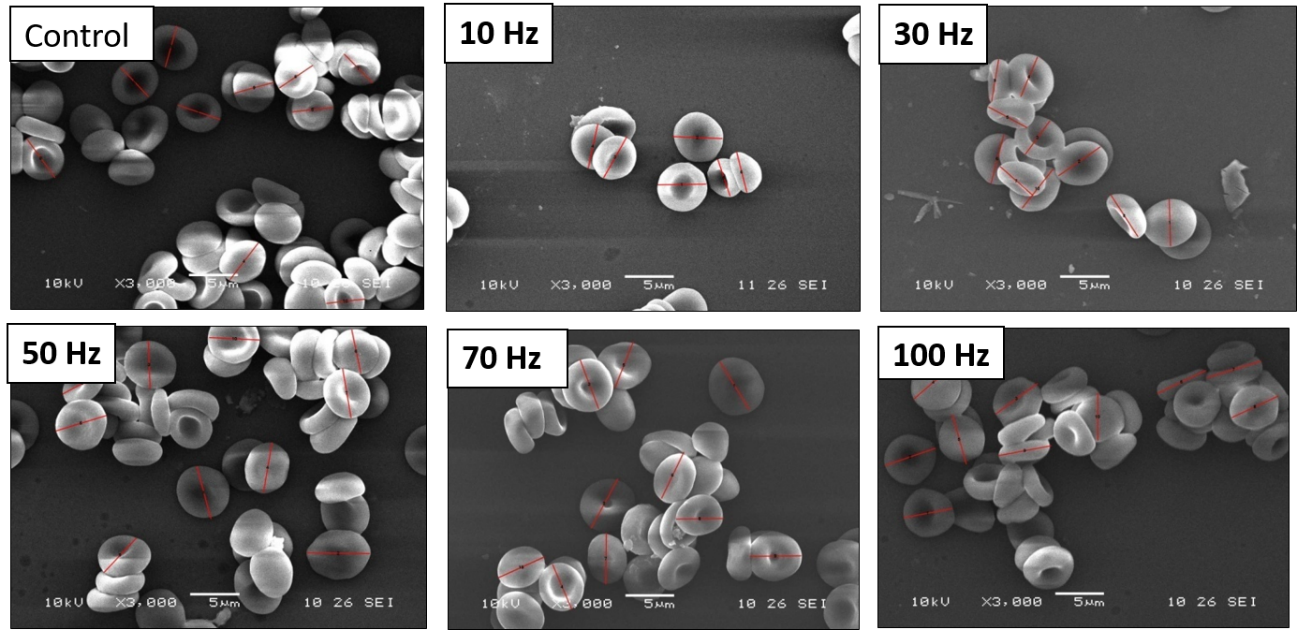

**Figure S2.** Morphology of isolated RBC from vibrated whole blood at different frequencies of vibrations i.e. 5 Hz, 10 Hz, 20 Hz, 30 Hz, 40 Hz, 50 Hz, 60 Hz, 70Hz and 100 Hz with constant amplitude

The reason for the decrease in thickness and increase in diameter of the RBC could be due to the impact of mechanical vibration, where mechanical stresses are developed at the cytoplasm of RBC.

**Experimental procedure for surface tension measurement**

The pendant drop method was used to study the effect of vibration on surface tension of RBC, Whole Blood, and blood plasma. The setup consists three parts an illuminating, viewing system to visualize the drop and a data acquisition system. The setup was calibrated by using 1 ml of distilled water taken in a syringe. The Young-Laplace equation, which links interfacial tension to drop shape to determine surface tension by using SCA20 software. Surface tension was measured before and after mechanical vibrations of whole blood, red blood cell and plasma samples.

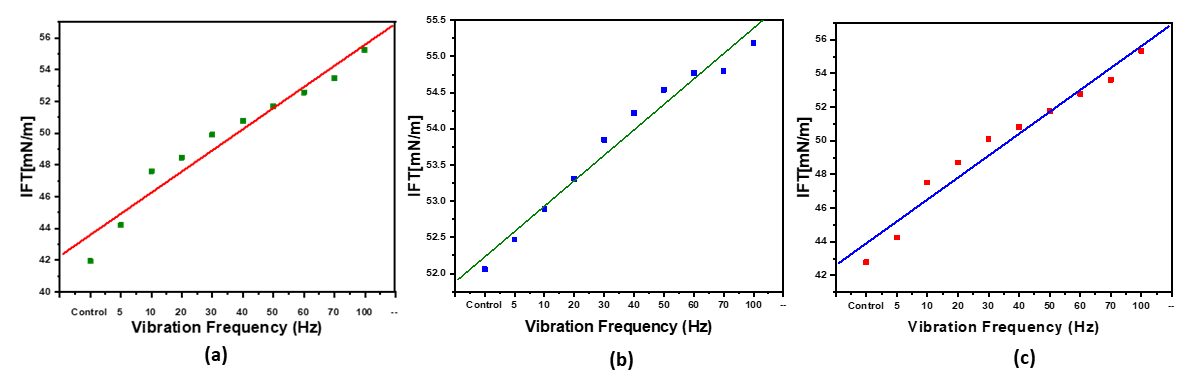
**Surface Tension of Whole Blood, RBC and plasma**

**Figure *S3.*** Surface tension of (a) whole blood (b) RBC (c) plasma at different vibration frequencies i.e. 5 Hz, 10 Hz, 20 Hz, 30 Hz, 40 Hz, 50 Hz, 60 Hz, 70Hz and 100 Hz with constant amplitude

Surface tension of blood and its components were measured by using pendant drop method.**^3^** Surface tension of whole blood shows increase from 41.95 mN/m to 55.23 mN/m with increasing vibrating frequencies **(Figure S3(a))**. The percentage difference in surface tension is 5 % for 5 Hz, 13 % for 10 Hz, 16 % for 20 Hz, 19 % for 30 Hz, 21 %, for 40 Hz, 23 % for 50 Hz, 25 % for 60 Hz, 27 % for 70 Hz, and 32 % for 100 Hz as compared to control.

Surface tension of RBC **(Figure S3(b))** shows increase from 42.78 mN/m to 55.33 mN/m with increasing vibrating frequencies. The percentage difference in surface tension is 3 % for 5 Hz, 11 % for 10 Hz, 14 % for 20 Hz, 17 % for 30 Hz, 19 % for 40 Hz, 21 % for 50 Hz, 23 % for 60 Hz, 25 % for 70 Hz, and 29 % for 100 Hz as compared to control.

Surface tension of plasma **(Figure S3(c))** shows increase from 52.06 mN/m to 55.19 mN/m with increasing vibrating frequencies. The percentage difference in surface tension is 1 % for 5 Hz, 2 % for 10 Hz, 2 % for 20 Hz, 3 % for 30 Hz 4 %, for 40 Hz, 5 % for 50 Hz, 5 % for 60 Hz, 5 % for 70 Hz, and 6 % for 100 Hz as compared to control.

The spectra of surface tension of whole blood, rbc and plamsa showed that the surface tension is directly proportional to the vibration frequency

**Experimental Procedure for measurement of viscosity**

Oswald viscosity meter was used for determining relative viscosity of blood samples. It measured the time required for the sample to flow through viscometer. The flow time is proportional to the viscosity of the sample. Nine ml of solvent sample (saline) was taken in Oswald viscosity meter tube and time required was measured (T_o_). Similar experimental procedure were repeated for the 9 ml of solute sample (vibrated blood) (T_1_). Relative viscosity of Human whole blood, Red Blood Cell and Plasma were measured using following formula:

Relative viscosity η_r_ = (T_1_ x ρ_1_)/(T_o_ x ρ_o_)

η_r_ = Relative viscosity

T_1_=Time of flow for solute (whole blood,Red Blood Cell and Plasma)

T_o_=Time of flow for solvent (saline water)

ρ_o_=Density of solvent

ρ_1_=Density of solute (blood/ RBC/Plasma+saline)

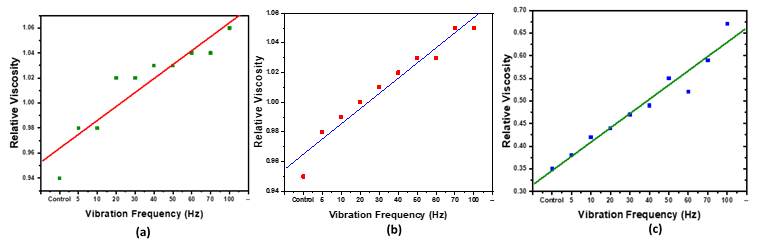

**Figure S4:**Relative viscosity of (a) whole blood (b) RBC (c) plamsa at different vibration frequencies i.e. 5 Hz, 10 Hz, 20 Hz, 30 Hz, 40 Hz, 50 Hz, 60 Hz, 70Hz and 100 Hz with constant amplitude

It was seen that the percentage difference in Relative viscosity in whole blood, RBC, and plasma in Figure S4.

**Experimental Procedure for Osmotic Fragility**

Osmotic fragility of RBC and whole blood were measured by taking the ratio of absorption of vibrated sample with the non vibrated sample. The ratio of surpernant of vibrated blood to absorbance of surpernant of control blood (non-vibrated) was measured at 540 nm wavelength.**^4^** The optical density of the vibrated and nonvibrated were measured with reference to the distilled water.

For osmotic fragility test the blood samples were prepared by taking NaCl solution of 0.45 % concentration which were prepared using 0.9 % saline. In each of the tube containing 10 ml of dilute NaCl solution, 0.05 ml of blood subjected to vibration was added. Both samples were mixed gently and allowed to stand for 30 minutes. If there was no hemolysis, the red cells were found at the bottom of the tubes with clear saline solution above. If some hemolysis occurred, the saline was tinged red with hemoglobin. If hemolysis was complete, the fluid was uniform in color throughout and there were no red cells visible at the bottom of the tube.

**Osmotic Fragility of RBC**

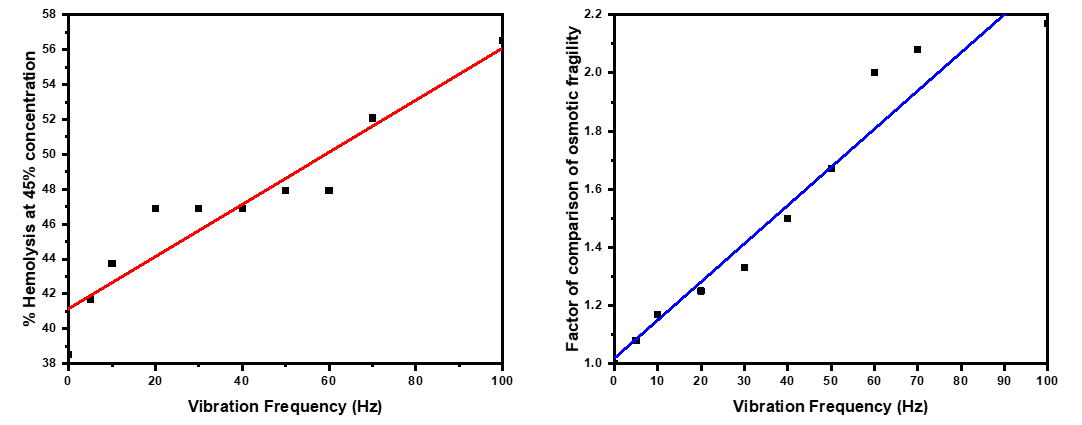

**(a)**

**(b)**

**Figure S5**(a) % Hemolysis at 45 % concentration and (b) Factor of comparison of osmotic fragility of whole blood (control and vibrated) at different frequencies i.e. 5 Hz, 10 Hz, 20 Hz, 30 Hz, 40 Hz, 50 Hz, 60 Hz, 70Hz and 100 Hz with constant amplitude

The percentage difference in % Hemolysis at 0.45 % concentration **(Figure S5(a))** of mechanically vibrated and non vibrated whole blood is 4 % for 5 Hz, 14 % for 10 Hz, 22 % for 20 Hz, 11 % for 30 Hz 17 %, for 40 Hz, 9 % for 50 Hz, 14 % for 60 Hz, 24 % for 70 Hz, and 47 % for 100 Hz as compared to control. The percentage difference in factor of comparison of osmotic fragility (Figure 13(b)) of mechanically vibrated and non vibrated whole blood is 8 % for 5 Hz, 17 % for 10 Hz, 25 % for 20 Hz, 33 % for 30 Hz 50 %, for 40 Hz, 67 % for 50 Hz, 100 % for 60 Hz, 108 % for 70 Hz, and 47 % for 117 Hz as compared to control.

**Theoretical Interpretation**

**Determining natural frequency of Red Blood Cell (RBC)**

Using Ansys software, the natural frequency of RBC was measured. Ansys software is a simulation software which generates physical stresses on materials usinf finite element analysis principles. It performs structural analysis of the material subjected to various types of stresses.

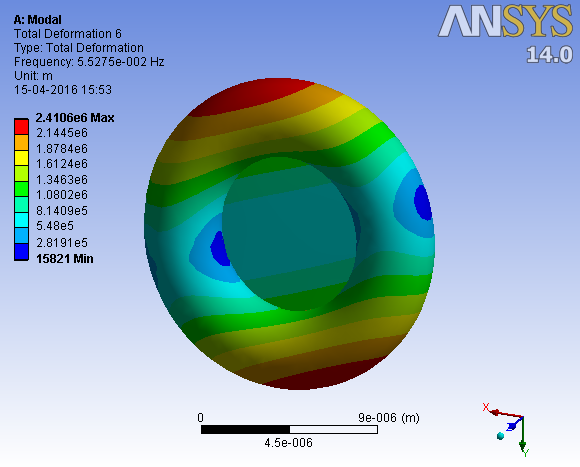

**Figure S6.** Natural frequency of RBC.

Physical parameters viz. shape, density of RBC is 1.125 g/cm^3^, Young’s modulus is 0.1 kPa, Poisson’s ratio is 0.5 and mass is 0.009864 g. These input parameters were fed to Ansys software. Deformation of RBC is observed at at natural frequency of 55.27 mHz **(Figure S6)**.

**Stresses developed on Red Blood cell due to mechanical vibrations**

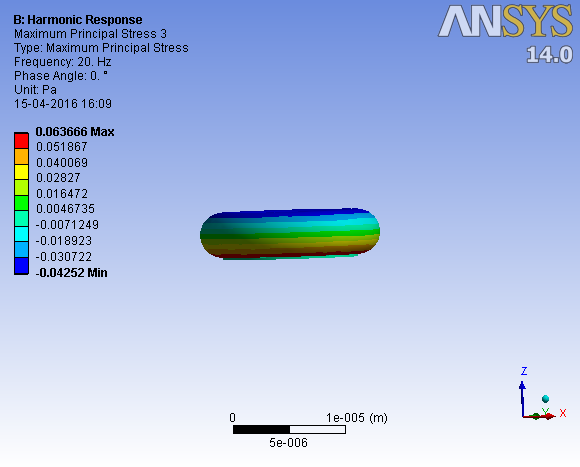

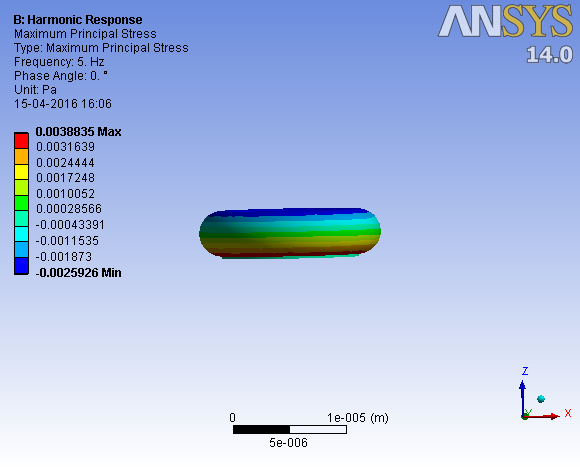

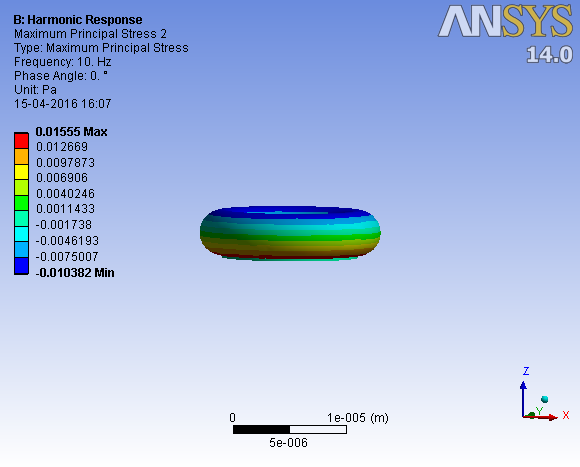

**(a)**

**(b)**

**(c)**

**Figure S7 (a-c).** Stresses profile on Red Blood Cell due to mechanical vibration at 5 Hz, 10 Hz and 15 Hz

Develoment of mechanical stresses in the region of cytoplasm of RBC is seen **(Figure S7(a-c)**.

Theis mechanical stress was observed by using Ansys software. Mechanical vibrations may affect the deformation of RBC permenentally which can lead to health problem. It is suggested to avoid exposure for continuous mechanical vibrations to the human body.

Representative data from one sample set of Diameter of RBC (control and vibrated) is given below.

|  | **Control** | **5 Hz** | **10Hz** | **20 Hz** | **30 Hz** | **40 Hz** | **50 Hz** | **60 Hz** | **70 Hz** | **100Hz** |
| --- | --- | --- | --- | --- | --- | --- | --- | --- | --- | --- |
| Label  RBC | D  (µm) | Dia  (µm) | Dia  (µm) | D µm) | D  (µm) | D (µm) | D (µm) | D (µm) | D  (µm) | D  (µm) |
| 1 | 4.86 | 4.38 | 4.92 | 4.66 | 5.46 | 5 | 5.46 | 5.82 | 5.84 | 5.59 |
| 2 | 5.17 | 5.14 | 5 | 4.68 | 4.49 | 5.45 | 5.62 | 5.68 | 5.63 | 5.87 |
| 3 | 4.65 | 5 | 4.81 | 4.99 | 5.15 | 5.04 | 5.5 | 5.49 | 5.17 | 5.49 |
| 4 | 5 | 4.74 | 4.73 | 4.64 | 5.03 | 5.68 | 5.16 | 5.4 | 5.44 | 5.95 |
| 5 | 4.56 | 5.14 | 5.41 | 5.15 | 5.49 | 5.07 | 5.35 | 5.54 | 5.05 | 5.75 |
| 6 | 4.33 | 4.63 | 4.99 | 5.12 | 5.56 | 4.88 | 5.83 | 4.49 | 5.7 | 5.52 |
| 7 | 4.68 | 5 | 4.85 | 4.99 | 4.74 | 5.73 | 4.94 | 5.27 | 5.83 | 5.86 |
| 8 | 4.68 | 4.3 | 5.03 | 5.06 | 5.2 | 4.89 | 4.62 | 4.82 | 5.01 | 5.46 |
| 9 | 4.96 | 4.7 | 4.74 | 5.14 | 5.13 | 4.75 | 4.81 | 5.22 | 5.06 | 5.53 |
| 10 | 4.72 | 5.17 | 4.71 | 5.55 | 4.95 | 5.63 | 5.23 | 5.86 | 5.6 | 5.41 |
| **Mean** | **4.76** | **4.82** | **4.92** | **5** | **5.12** | **5.21** | **5.25** | **5.36** | **5.43** | **5.64** |
| **Std. Dev** | **0.24** | **0.32** | **0.21** | **0.28** | **0.34** | **0.37** | **0.38** | **0.43** | **0.33** | **0.2** |

The output data of elongation index from SEM images as given in below table.

| **Control** | | | **10 Hz** | | | **30 Hz** | | | **50 Hz** | | |
| --- | --- | --- | --- | --- | --- | --- | --- | --- | --- | --- | --- |
| L | W | EI | L | W | EI | L | W | EI | L | W | EI |
| 5.32; 5.37; 5.14; 4.59; 5.37; 5.21; 4.88 | 5.18; 4.46; 4.3; 4.53; 4.96; 4.77; 4.81 | 0.013; 0.092; 0.088; 0.006; 0.039; 0.044; 0.007 | 5.28; 5.31; 4.91; 5.35; 5.09; 5.31; **5.11** | 4.64; 4.88; 4.63; 4.32; 4.8; 4.32; 4.86 | 0.064; 0.042; 0.029; 0.106; 0.029; 0.102; 0.025 | 5.09; 5.64; 5.23; 5.83; 5.18; 5.55; 5.43 | 4.74; 5.27; 5.15; 4.91; 4.83; 4.91; 4.3 | 0.035; 0.033; 0.007; 0.085; 0.034; 0.061; 0.116 | 5.68; 5.79; 6.59; 5.44; 6.22; 5.76; 6.23 | 5.11; 5.22; 5.49; 5.10; 4.30; 5.55; 5.59 | 0.052; 0.051; 0.091; 0.032; 0.182; 0.018; 0.054 |
| **Mean** | | **0.038** |  |  | **0.057** |  |  | **0.053** |  |  | **0.069** |

| **70 Hz** | | | **100 Hz** | | |
| --- | --- | --- | --- | --- | --- |
| L | W | EI | L | W | EI |
| 6.04; 6.96; 5.32; 6.77; 6.05; 5.96 | 4.98 \|; 5.97 \|; 4.54 \|; 5.68 \|; 5.77 \|; 4.08 \| | 0.096 \|; 0.076 \|; 0.079 \|; 0.087 \|; 0.023 \|; 0.187 \| | 6.25; 6.24; 5.28; 5.8; 5.86; 6.14 | 5.00; 5.01; 5.2; 4.39; 4.82; 4.92 | 0.111; 0.109; 0.007; 0.138; 0.097 0.110 |
| **Mean** | | **0.092** |  | | **0.101** |

**L: Length of major axis of RBC; W: Length of minor axis; EI: Elongation Index**

**References:**

1. Atkins, C. G., Buckley, K., Blades, M. W., & Turner, R. F.

Raman spectroscopy of blood and blood components.

Applied spectroscopy, 71(5), (2017), 767-793.

2. Alummoottil, S., van Rooy, M. J., Bester, J., Grobbelaar, C., & Phulukdaree, A.

Scanning Electron and Atomic Force Microscopic Analysis of Erythrocytes in a Cohort of Atopic Asthma Patients—A Pilot Study.

Hemato, 4(1), (2023), 90-99.

3. Yadav, S. S., Sikarwar, B. S., Ranjan, P., Janardhanan, R., & Goyal, A.

Surface tension measurement of normal human blood samples by pendant drop method.

Journal of Medical Engineering & Technology, 44(5), (2020), 27-236.

4. Walski, T., Chludzińska, L., Komorowska, M., & Witkiewicz, W.

Individual osmotic fragility distribution: a new parameter for determination of the osmotic properties of human red blood cells.

BioMed Research International, 2014(1), 162102.
